# Peripheral nerve injury induces an immunometabolic signature involving reduced free fatty-acid pools

**DOI:** 10.64898/2026.01.13.699159

**Authors:** Jun Seo Park, Hye Won Jun, Jaesung Lee, Sung Joong Lee

## Abstract

Peripheral nerve injury-induced persistent pain hypersensitivity cannot be fully explained by increased neuronal excitability alone. Rather, it is increasingly recognized as a pathological tissue state characterized by sustained glial activation, immune signaling, and synaptic remodeling within the spinal cord. In this context, metabolic remodeling of the spinal cord has emerged as a potentially important feature of neuropathic pain. However, the metabolic patterns associated with this state, and their relationship to underlying molecular programs, remain incompletely defined. Here, we performed GC–MS-based untargeted metabolomic profiling of the spinal cord dorsal horn on day 7 after spinal nerve transection (SNT). To provide orthogonal validation, we integrated pathway-analysis results from four independent spinal cord RNA-sequencing datasets derived from distinct neuropathic pain models and further conducted qPCR-based validation. Metabolic profiling revealed a clear separation between SNT and sham samples, marked by broad depletion of the free fatty acid pool and features consistent with an immunometabolic shift. Consistently, analyses across RNA-sequencing datasets and qPCR validation demonstrated upregulation of immune and inflammatory programs, together with downregulation of fatty acid metabolism and cholesterol homeostasis. Collectively, these findings suggest that persistent neuropathic pain should be interpreted not simply as a consequence of neuronal and immune signaling, but also through the metabolic tissue environment that supports and sustains this pathological state.

## INTRODUCTION

Neuropathic pain is a disorder characterized by persistent pain hypersensitivity that arises following nerve injury^1^. Although peripheral nerve damage initiates the pathological cascade, the transition to chronic pain requires sustained adaptations within the central nervous system, including the spinal cord^2,3^. In particular, spinal nociceptive circuits undergo persistent sensitization accompanied by glial activation and inflammatory signaling, and these processes collectively contribute to the maintenance of mechanical hypersensitivity^3-8^.

Beyond cytokine- and synapse-centered mechanisms, accumulating evidence indicates that chronic neuroinflammation is coupled to metabolic reprogramming^9-13^. This immunometabolic state is characterized by alterations in central carbon metabolism, redox homeostasis, and lipid utilization, and is increasingly recognized as a fundamental determinant of immune-cell activation and persistence^14-15^. During neuropathic pain, the spinal cord can harbor both resident glial populations and infiltrating immune cells, and metabolic rewiring in this context may shape both inflammatory outputs and neuronal excitability^4,7,9,11^. Indeed, recent studies have begun to report changes in metabolic patterns during pain chronification across central nervous system regions, including the spinal cord^9-13^. From this perspective, regulation of specific pathways—such as lactate production and glycolysis—has been proposed to contribute to the development and maintenance of chronic pain^11^. However, several important gaps remain: untargeted metabolomic profiling of the spinal cord under established chronic neuropathic pain states is still limited; consequently, (i) an unbiased metabolomic landscape of the spinal cord during persistent pain hypersensitivity has not been sufficiently delineated, and (ii) major metabolic nodes that reproducibly track with inflammatory programs across neuropathic pain models have not been fully defined.

Here, we performed gas chromatography–mass spectrometry (GC–MS)-based untargeted metabolomics of spinal cord tissue at a time point of sustained pain hypersensitivity, when glial activation was most pronounced. Through clustering and pathway-level analyses, we comprehensively characterized metabolomic remodeling in the spinal cord during chronic neuropathic pain. To further test whether these metabolic remodeling signatures generalize across paradigms, we additionally conducted a meta-analysis of four independent spinal cord RNA-seq datasets derived from distinct chronic pain models. For orthogonal validation, we further assessed the expression of key metabolic nodes by quantitative PCR (qPCR). Collectively, our data support an immunometabolic shift in the spinal cord during chronic pain, marked by increased inflammatory metabolism alongside depletion of the free fatty acid pool. These findings advance mechanistic understanding of pain chronification and highlight metabolic remodeling as a potentially actionable axis for therapeutic intervention.

## RESULTS

### Peripheral nerve injury is accompanied by long-term pain hypersensitivity and increased inflammatory markers

To model long-lasting neuropathic pain, we induced peripheral nerve injury by L5 spinal nerve transection (SNT; Fig. 1A)^16-18^. Mechanical sensitivity was monitored over 7 days post-surgery using the von Frey assay (Fig. 1B). Relative to sham-operated controls, SNT mice exhibited sustained mechanical hypersensitivity across the observation window, consistent with the establishment of a long-lasting hypersensitivity state.

**Figure 1.**
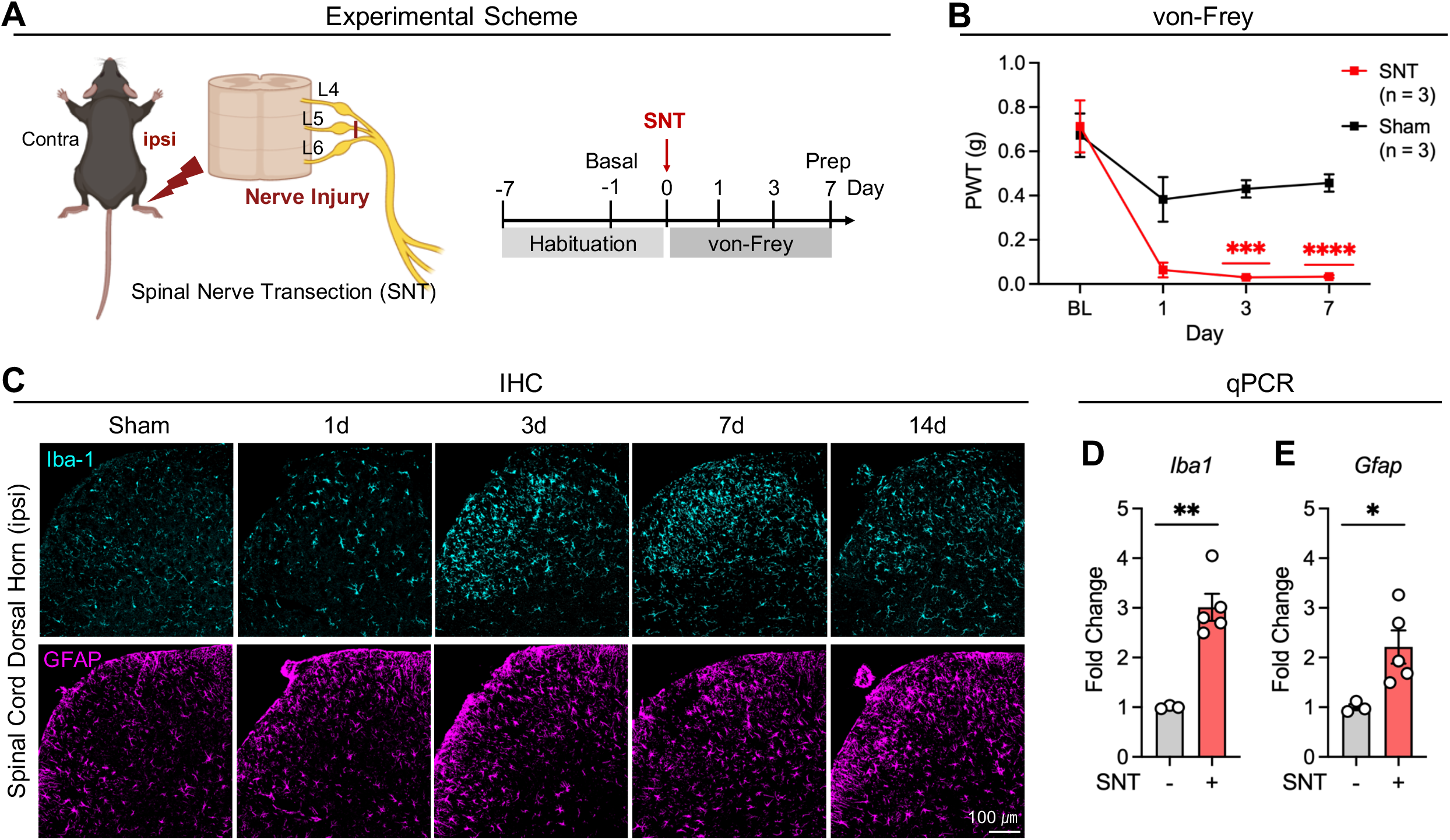
Peripheral nerve injury is accompanied by long-term pain hypersensitivity and increased inflammatory markers. (A) Schematic illustration of the spinal nerve transection (SNT) model and experimental timeline. Following habituation and baseline assessment, SNT was performed at day 0, and mechanical sensitivity was evaluated by von Frey testing at the indicated time points. (B) von Frey test showing paw withdrawal threshold (PWT) in SNT and sham groups across baseline (BL) and post-operative days 1, 3, and 7 (n = 3 per group). ***P = 0.0002 (3 day SNT vs 3 day Sham), ****P < 0.0001 (7 day SNT vs 7 day Sham) (C) Representative confocal images of Spinal Cord (Dorsal Horn ipsi area) (Scale bar, 100 μm). IHC smple of GFAP positive and Iba1 positive cells. (D, E) Spinal cord qPCR data at day 7 time point. (SNT; n = 5, Sham; n = 3) (F) Fold change of *Iba1*. **P = 0.0017 (SNT vs Sham). (E) Fold change of *Gfap*. *P = 0.0208 (SNT vs Sham). Data are represented as the mean□±□SEM; \**P*□ < 0.05, **P < 0.01, \*\*\**P*□ <□0.001, \*\*\*\**P* □<□0.0001; Two-way ANOVA followed by Bonferroni’s multiple comparisons test (B); Student’s t-test (D, E).

We next examined temporal changes in glial activation markers during persistent pain hypersensitivity. Using immunohistochemistry (IHC)-based histological analysis and qPCR-based gene-expression analysis, we assessed ionized calcium-binding adaptor molecule 1 (Iba1), a microglial marker, and glial fibrillary acidic protein (GFAP), a marker of reactive astrocytes. Histological analysis revealed a progressive increase in glia-associated inflammatory markers following nerve injury, with elevated levels persisting through day 7 (Fig. 1C). Consistently, gene-expression analysis showed a gradual increase in inflammatory markers up to day 7 (Fig. S1), including significant upregulation of Iba1 and Gfap in the ipsilateral spinal cord at this time point (Fig. 1D, E).

Previous studies have reported key physiological alterations at day 7 in neuropathic pain, including astrocytic and microglial activation, inflammation, altered neuronal activity, and synaptic remodeling^17-20^. On the basis of these observations, we sought to characterize metabolic profile changes at the day 7 time point.

### Untargeted GC–MS metabolomics reveals a global shift in the spinal cord metabolomic landscape

To define pain-associated metabolic alterations in the spinal cord, we collected ipsilateral lateral L3–L5 spinal cord tissue at day 7 post-surgery and performed GC–MS–based untargeted metabolomics (Fig. 2A). Samples were processed through homogenization, heat-based enzyme inactivation, and centrifugation prior to GC–MS analysis (Fig. 2A). We first assessed whether neuropathic pain was associated with an overall shift in metabolomic profiles. Principal component analysis (PCA) separated SNT and sham samples (Fig. 2B), and supervised partial least squares-discriminant analysis (PLS-DA) showed a concordant separation pattern (Fig. 2C). The statistical reliability of clustering in each analysis was confirmed (Fig. S2). Notably, metabolites aligned with the SNT direction included lactate, itaconate, and glycine, whereas sham samples aligned with lipid-associated metabolites such as oleate/linoleate/vaccenate in the loading structure of the multivariate models (Fig. 2B–C). Together, these analyses indicate that neuropathic pain is accompanied by a robust global remodeling of the spinal cord metabolomic state at the chronic time point.

**Figure 2.**
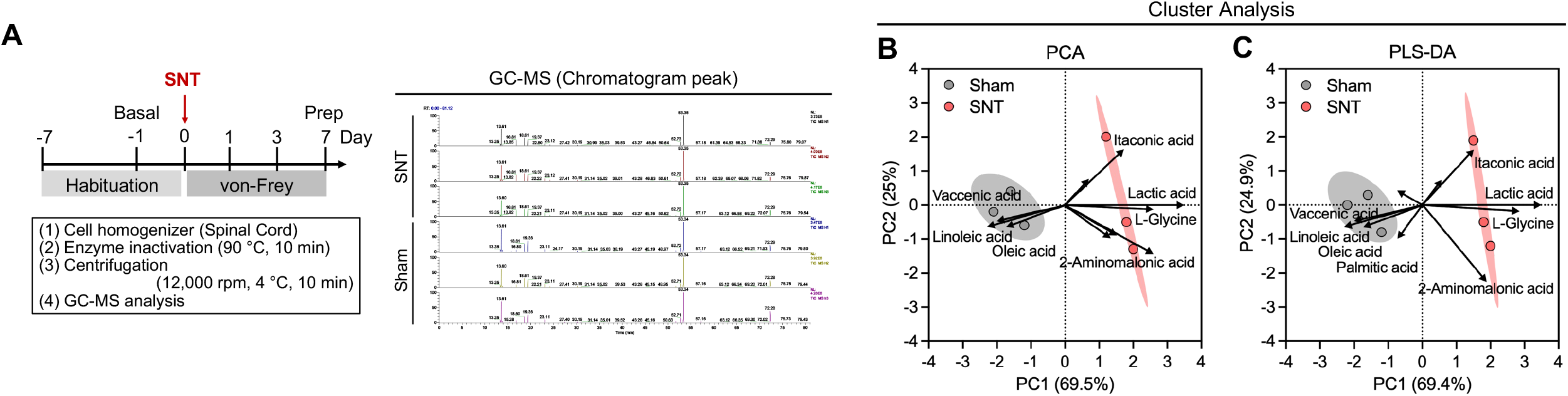
GC–MS-based metabolomics distinguishes SNT from sham spinal cord at the chronic phase. (A) Schematic illustration of spinal cord metabolomics at day 7 post-SNT, including tissue processing steps and representative GC–MS chromatogram peaks from sham and SNT samples. (SNT; n = 3, Sham; n = 3) (B and C) Cluster analysis of GC-MS data; compared by groups (sham, SNT). Show axis of representative metabolite. Arrows indicate loading vectors for representative metabolites (lactate, itaconic acid, glycine, and fatty acids). (B) Principal component analysis (PCA) of GC-MS fold-change profiles of each groups. Cluster separation was observed in SNT vs sham groups. The variance in X-axis (PC1) is 69.5% and on Y-axis (PC2) is 25%. (C) Partial least squares-discriminant analysis (PLS-DA) of GC-MS fold-change profiles of each groups. Cluster separation was observed in SNT vs sham groups. The variance in X-axis (PC1) is 69.4% and on Y-axis (PC2) is 24.9%.

### Sustained pain hypersensitivity is characterized by immunometabolic reprogramming and concomitant depletion of the fatty acid pools

We next quantified individual metabolite alterations in SNT relative to sham controls. Across the metabolite panel, SNT samples showed a coordinated pattern in which subsets of organic acids and amino acids increased, while fatty acid–related metabolites broadly decreased (Fig. 3A–B). Among the most prominent increases, lactate, itaconate, and L-glycine were significantly elevated in SNT compared with sham (Fig. 3C–E). In contrast, multiple lipid-related metabolites—including palmitate, stearate, oleate, linoleate, and vaccenate—were significantly reduced in SNT (Fig. 3F–J). This bidirectional signature (selected organic acids/amino acids up; broad fatty-acid pool down) was consistently captured by fold-change and significance-based prioritization in the volcano-plot summary (Fig. 3K). To integrate metabolite-level changes at the pathway scale, we performed metabolite set enrichment analysis (MSEA)^21-22^. SNT samples showed enrichment of pathways linked to amino-acid and short-chain organic-acid metabolism (e.g., arginine/proline-related and butanoate-related modules; Fig. 3L), whereas lipid-associated pathways—including fatty-acid elongation and related lipid metabolic programs—were comparatively suppressed (Fig. 3M). Notably, energy metabolism–related modules, including the TCA cycle, also appeared among the pathways with decreased enrichment signals in this analysis (Fig. 3M). Collectively, these metabolomics data define a long-lasting neuropathic pain–associated spinal cord signature characterized by an immunometabolism-linked metabolite axis (lactate/itaconate/glycine) together with coordinated depletion of fatty-acid pools.

**Figure 3.**
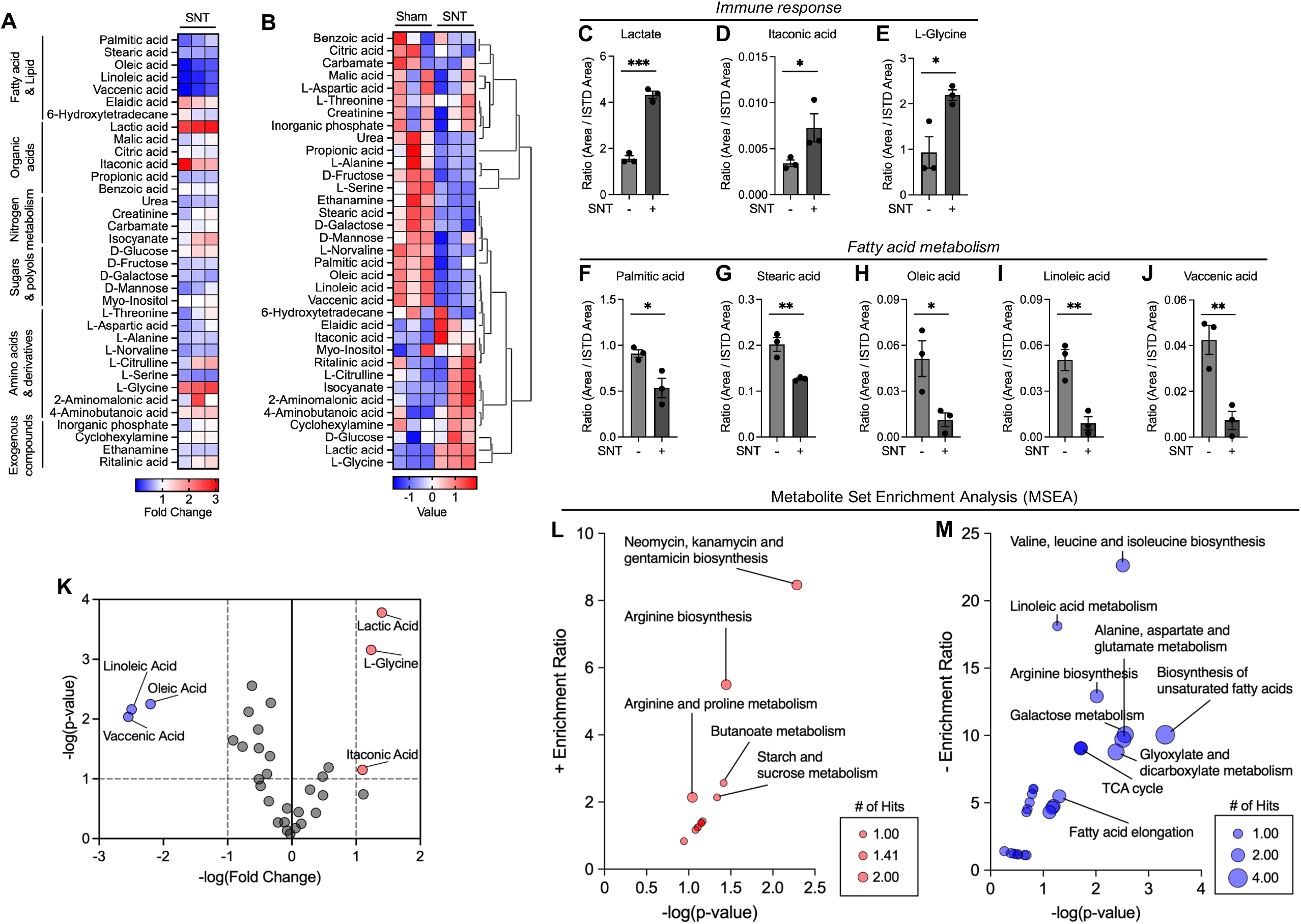
Chronic neuropathic pain is associated with an immunometabolic signature and depletion of fatty-acid pools. (A) Heatmap summarizing metabolite-class–level changes (fold change) in SNT relative to sham across major metabolite categories. (B) Heatmap of relative metabolite abundances across individual samples with hierarchical clustering. (C–E) Increased levels of immunometabolism-associated metabolites (lactate, itaconic acid, and L-glycine) in SNT compared with sham. (C) Relative concentration of lactate. ***P = 0.0002 (Sham vs SNT) (D) Relative concentration of itaconic acid. *P = 0.0250 (Sham vs SNT) (E) Relative concentration of L-glycine. *P = 0.0257 (Sham vs SNT) (F–J) Decreased levels of representative fatty acids (palmitic acid, stearic acid, oleic acid, linoleic acid, and vaccenic acid) in SNT compared with sham. (F) Relative concentration of palmitic acid. *P = 0.0291 (Sham vs SNT) (G) Relative concentration of stearic acid. **P = 0.0077 (Sham vs SNT) (H) Relative concentration of oleic acid. *P = 0.0330 (Sham vs SNT) (I) Relative concentration of linoleic acid. **P = 0.0071 (Sham vs SNT) (J) Relative concentration of vaccenic acid. **P = 0.0092 (Sham vs SNT) (K) Volcano plot depicting differential metabolites between SNT and sham, with representative up- and downregulated metabolites annotated. (L-M) Metabolite set enrichment analysis (MSEA) of GC-MS data; compared by each groups (Sham vs SNT). Labeling selected high enrichment metabolic pathway. Bubble size reflects the number of metabolite hits, and axes denote enrichment ratio and statistical significance (−log(p-value)). (L) Pathways enriched among increased metabolites (red). (M) Pathways enriched among decreased metabolites (blue). Data are represented as the mean□±□SEM; \**P*□ < 0.05, **P < 0.01, \*\*\**P*□ <□0.001, \*\*\*\**P* □<□0.0001; Student’s t-test (C-J).

### Transcriptomic analysis across neuropathic pain models reproduces immune activation and suppression of lipid metabolic programs

We next asked whether this metabolite signature was reflected at the transcriptional level across independent neuropathic pain paradigms. We re-analyzed four publicly available spinal cord RNA-seq datasets spanning spared nerve injury (SNI)^23^, chronic constriction injury (CCI; two independent datasets)^24-25^, and paclitaxel-induced peripheral neuropathy (PIPN)^26^, sampled at post-injury time points corresponding to established mechanical hypersensitivity (day 7 or day 14; Fig. 4A). Across datasets, gene set enrichment analysis (GSEA) consistently showed enrichment of inflammatory gene programs, exemplified by the “inflammatory response” signature (Fig. 4B–E). In contrast, lipid-associated metabolic programs were consistently suppressed, exemplified by depletion of the “fatty acid metabolism” gene set across models (Fig. 4F–I). Summary NES patterns across Hallmark and KEGG collections further highlighted a convergent directionality: immune-associated pathways were upregulated, whereas lipid metabolic pathways were downregulated in all four models (Fig. 4J–K, Fig. S3, Fig S4, Fig. S5, Fig. S6).

**Figure 4.**
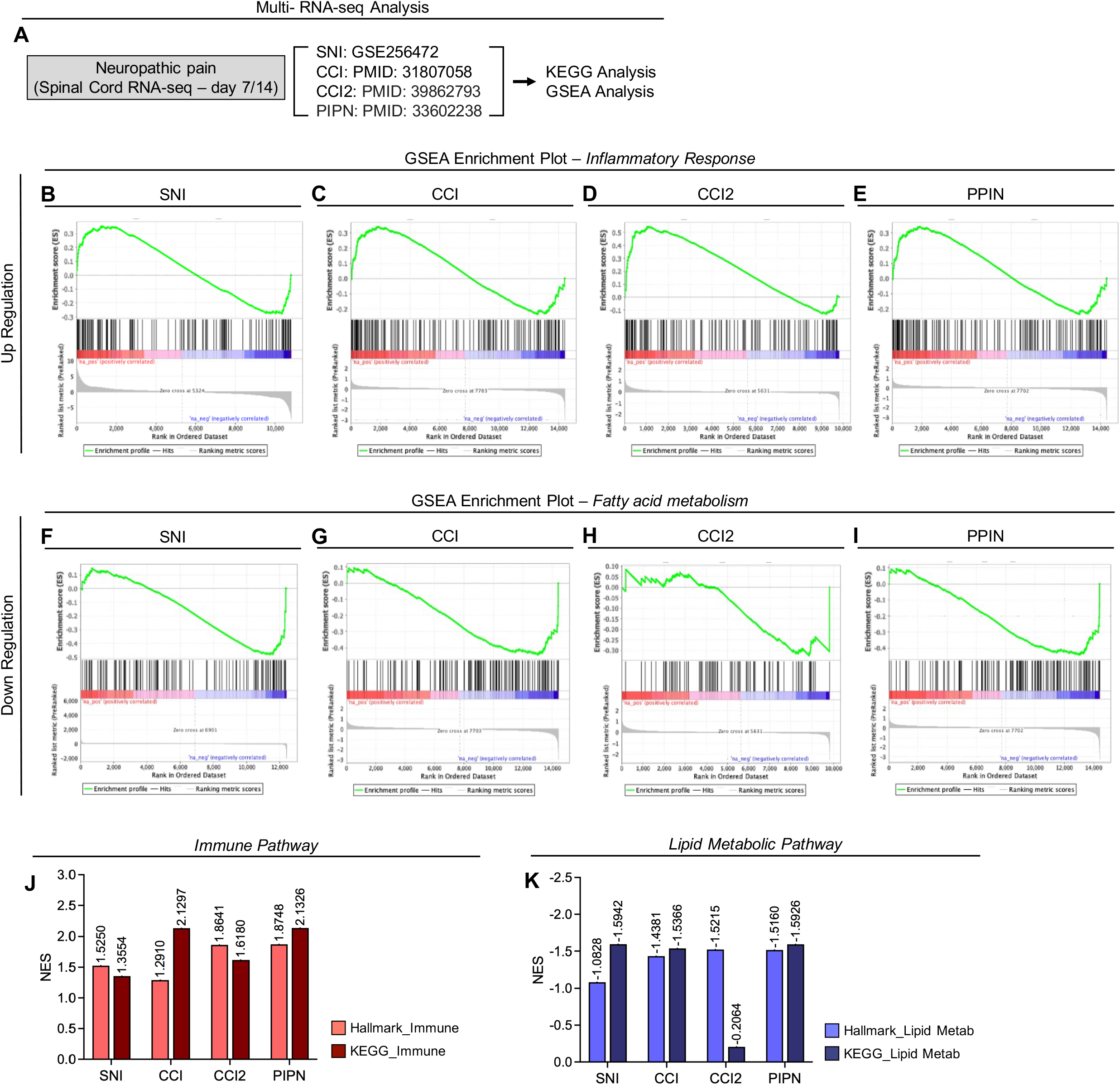
Cross-model spinal cord RNA-seq analyses reveal conserved inflammatory upregulation and lipid metabolic suppression. (A) Overview of the RNA-seq analysis framework using four independent spinal cord neuropathic pain datasets (SNI: GSE256472; CCI: PMID 31807058; CCI2: PMID 39862793; PIPN: PMID 33602238) sampled at day 7/14, followed by Hallmark GSEA and KEGG pathway analysis. (B–E) Representative GSEA enrichment plots showing consistent upregulation of an inflammatory response signature across models (SNI, CCI, CCI2, and PIPN). (F–I) Representative GSEA enrichment plots showing consistent downregulation of fatty acid metabolism across models. Across datasets, comparing Hallmark and KEGG pathway-level results. (J–K) Summary of normalized enrichment scores (NES) (J) Immune-related pathways; Immune programmes were defined by enrichment of Hallmark immune signatures (for example, Inflammatory response, with additional enrichment of IL6–JAK–STAT3 signalling, Interferon-γ response and Allograft rejection) and by KEGG immune-associated terms capturing cytokine signalling and innate immune effector modules (including Cytokine–cytokine receptor interaction, JAK–STAT signalling, Complement and coagulation cascades, Phagosome, Leukocyte transendothelial migration and Neutrophil extracellular trap formation). (K) Lipid metabolic pathways; Lipid metabolic programmes were defined by enrichment of Hallmark metabolic signatures (for example, Fatty acid metabolism, with additional enrichment of Cholesterol homeostasis and Oxidative phosphorylation) and by KEGG lipid-related terms (including Fatty acid elongation, Steroid biosynthesis, Bile secretion and Glycosphingolipid biosynthesis).

Beyond pathway-level alterations, we sought to determine whether these changes were reflected in specific regulatory genes. We first examined the mean fold change of individual genes across the four spinal cord RNA-sequencing datasets described above. Genes involved in fatty acid metabolism, selected on the basis of the observed depletion of the fatty acid pool, showed a broadly reduced expression pattern under neuropathic pain conditions (Fig. 5A). Conversely, key immune-response genes associated with the metabolite alterations identified above exhibited an overall increase in expression (Fig. 5A).

**Figure 5.**
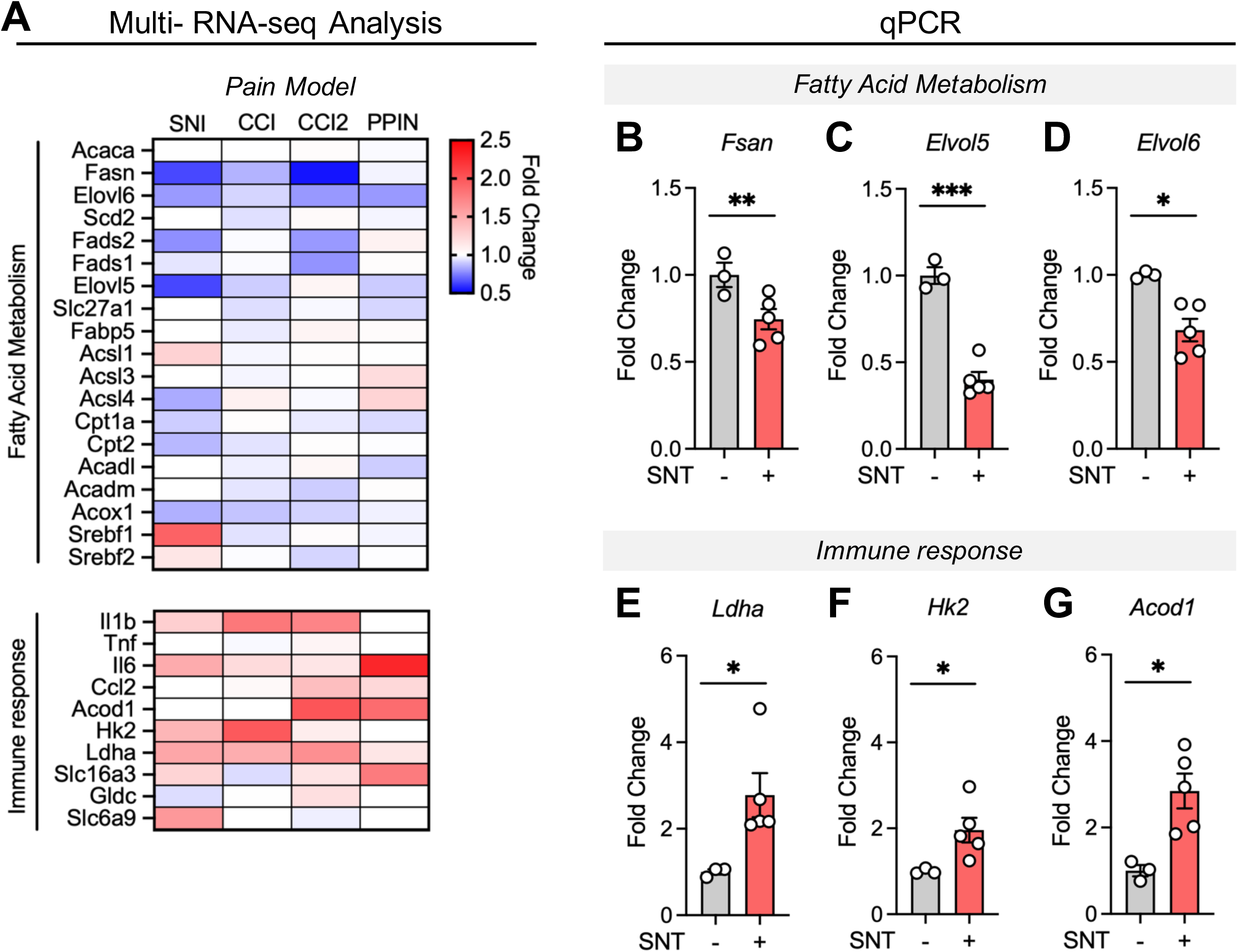
Targeted analysis and qPCR validation of lipid- and immune-associated transcriptional changes. (A) Heatmap of selected gene in each pain model RNA-seq data. Averge fold change of selected lipid metabolism-related and immune response-related genes. (B-G) Spinal cord qPCR data. (SNT; n = 5, Sham; n = 3) (B) Fold change of *Fsan*. **P = 0.0077 (SNT vs Sham). (C) Fold change of *Elvol5*. ***P = 0.0003 (SNT vs Sham). (D) Fold change of *Elvol6*. *P = 0.0107 (SNT vs Sham). (E) Fold change of *Ldha*. *P = 0.0404 (SNT vs Sham). (F) Fold change of *Hk2*. *P = 0.0474 (SNT vs Sham). (G) Fold change of *Acod1*. *P = 0.0148 (SNT vs Sham). Data are represented as the mean□±□SEM; \**P*□ < 0.05, **P < 0.01, \*\*\**P*□ <□0.001, \*\*\*\**P* □<□0.0001; Student’s t-test (B-G).

To orthogonally validate these findings in the SNT model, we performed qPCR analysis of spinal cord tissue collected 7 days after nerve injury. Genes involved in fatty acid metabolism and linked to the regulation of key metabolites identified by metabolomics—including fatty acid synthase (*Fasn*), elongation of very long-chain fatty acids protein 5 (*Elovl5*), and elongation of very long-chain fatty acids protein 6 (*Elovl6*)^27,28^—were significantly downregulated in the SNT group (Fig. 5B–D). By contrast, genes associated with immune-response-related metabolites, including lactate dehydrogenase A (*Ldha*), hexokinase 2 (*Hk2*), and aconitate decarboxylase 1 (*Acod1*)^29.30^, were significantly upregulated (Fig. 5E–G).

Taken together, transcriptomic analysis and orthogonal validation of key genes independently reproduced the directionality observed in our metabolomic dataset: reinforcement of immune-associated programs and attenuation of lipid metabolic programs. These findings support a coordinated depletion of fatty acid pools and a shift toward an immunometabolic state across the spinal cord during long-lasting neuropathic pain (Fig. 6).

**Figure 6.** Schematic Illustrations. Spinal Cord immunometabolic signature and lipid metabolism suppression of chronic neuropathic pain states. (Created in https://www.biorender.com)

## DISCUSSION

In this study, we present an untargeted metabolomic analysis of spinal cord tissue under long-lasting neuropathic pain conditions and delineate a coordinated shift in spinal metabolic signatures. We show that this pattern—strengthening of glycolytic/organic acid metabolism and immunometabolism-associated signatures alongside suppression of lipid metabolism—is not confined to the metabolite level, but is concordantly observed at the transcriptomic level and shared across models. These findings provide an unbiased framework in which chronic neuropathic pain is sustained, at the spinal cord level, through a coupled reorganization of immune and metabolic programs.

During the chronic phase, the spinal cord was characterized by elevation of a lactic acid–itaconic acid–glycine signature together with a broad and consistent depletion of major fatty-acid pools, including oleic acid and linoleic acid. Increased lactic acid is consistent with a glycolytic bias and associated microenvironmental changes within a chronically inflamed milieu, a mechanism increasingly implicated in glia-associated pain biology^11-12, 31^. The concomitant increases in itaconic acid and glycine are also consistent with immunometabolic reprogramming linked to stress-adaptive pathways^32-35^. Importantly, a central pathophysiological feature of our dataset is not fluctuation of a single lipid species, but rather broad depletion of multiple GC–MS-detectable fatty-acid pools spanning both saturated and unsaturated species^36^. This raises the possibility that, in the chronic pain spinal cord, reduced fatty-acid pools may reflect altered mobilization/uptake/esterification and/or sustained suppression of lipid metabolic programs, potentially accompanied by increased demand for membrane remodeling and signaling lipids^37^. Such changes imply a reprioritization of energy utilization and biosynthetic allocation in the chronic spinal cord, potentially creating a biochemical substrate that favors the long-term persistence of inflammatory outputs and altered neuronal excitability^38^. However, lipid alterations require a more nuanced interpretation than a simple global reduction. Fatty acid metabolism may be regulated in context-dependent and directionally distinct ways, and neuropathic pain may involve more complex lipid-associated changes, including pathological lipid imbalance arising from lipid remodeling and alterations in phospholipid composition^39,40^.

Our study further underscores that this coupled metabolic–immune signature is unlikely to be specific to a particular experimental condition or model. Across multiple independent spinal cord RNA-seq datasets derived from distinct neuropathic pain models, upregulation of immune-related programs and downregulation of lipid metabolic programs were consistently reproduced. This convergence supports the existence of a shared molecular “directionality” toward which the spinal cord may evolve under chronic pain. In doing so, these data provide evidence that higher-order biological features defining the chronic spinal state may be generalizable across paradigms—addressing, at least in part, longstanding concerns regarding reproducibility and cross-model generalization in pain research. Consequently, our work proposes a unifying layer of explanation that extends beyond neurotransmission and inflammatory signaling alone, positioning metabolic program remodeling as a fundamental and integrative dimension of pain chronification.

These findings also offer a conceptual foothold for therapeutic strategy. Chronic pain treatments have traditionally focused on modulating neuronal excitability or blocking specific inflammatory pathways; however, such approaches often fail to dismantle the persistence of the pathological state^41,42^. The immunometabolic transition and lipid metabolic suppression described here suggest that metabolic state may contribute to stabilization of the long-lasting pain state within the spinal cord, motivating an expansion of therapeutic hypotheses toward modulation of metabolic state^42-44^. In this view, metabolic remodeling is not a secondary epiphenomenon, but may constitute a regulatory axis that contributes to long-term maintenance of inflammation and circuit sensitization^38,45^. Accordingly, our results motivate testing whether interventions that modulate metabolic state can move beyond symptom suppression and attenuate the persistence of the pathological state.

### Limitations of the study

First, this study focused on a single post-injury time point, day 7 after SNT, and therefore does not define the temporal sequence through which the observed immunometabolic profile emerges or stabilizes. Second, the metabolomic analysis was performed with a limited sample size, which may constrain the robustness and generalizability of the metabolite-level findings. GC–MS-based profiling also captures a defined subset of detectable metabolites, including free fatty acids, and does not provide comprehensive lipidomic coverage or direct evidence of lipid metabolic flux. Thus, the observed reduction in free fatty-acid pools should not be interpreted as evidence for global lipid metabolic suppression. Third, because the analyses were performed using bulk spinal cord tissue, the cellular origins of these metabolic and transcriptional changes remain unresolved. Finally, although public RNA-seq re-analysis, qPCR, and IHC provide orthogonal support for the observed directionality, the present study does not establish causal links between metabolic remodeling, glial activation, and pain-like behavior. Future longitudinal, cell type-resolved, and perturbation-based studies will be required to define the mechanistic and functional relevance of this spinal immunometabolic signature.

## RESOURCE AVAILABILITY

### Lead contact

Further information and requests for resources and reagents should be directed to and will be fulfilled by the lead contact, Prof. Sung Joong Lee.

### Materials availability

All mouse lines and materials used in this study were provided or purchased from the mentioned companies or researchers. This study did not generate any new or unique reagents.

### Data and code availability

All data reported in this paper will be shared by the lead contact upon request.

This paper does not report original code.

All data associated with this study are present in the paper or the supplemental information.

## ACKNOWLEDGMENTS

We are grateful to all members of the Neuron-Glia Network Research Laboratory for their helpful discussions and help. This research was supported by the National Research Foundation of Korea (RS-2024-00402116; RS-2025-02215169 to S.J.L.).

## AUTHOR CONTRIBUTIONS

J.S.P. designed the research, performed most experiments, analyzed the data, and wrote the first draft of the manuscript. H.W.J. assisted with experimental procedures. J. L. provide validations. S.J.L. supervised the project and wrote the manuscript.

## DECLARATION OF INTERESTS

The authors declare no competing interests.

## METHODS

### Animals

The Institutional Animal Care and Use Committee of Seoul National University approved all animal experiments (Approval No. SNU-250905-4). All mice used were of the C57BL/6 strain. Male C57BL/6 mice (8–12□weeks of age) were purchased from DooYeol Biotech (Seoul, Korea). All animals were acclimated to standard conditions with a 12-hour light/dark cycle in a specific pathogen-free environment and given access to food and water *ad libitum*. All protocols were performed in accordance with guidelines from the International Association for the Study of Pain.

### Neuropathic pain mouse model

To generate a persistent pain model, a right L5 SNT was performed as previously described^12,18, 20^. Briefly, animals were anesthetized with isoflurane in an O_2_ carrier (induction 2% and maintenance 1.5%), and a small incision was made to expose the L4 and L5 spinal nerves. The L5 spinal nerve was then transected.

### Pain behavior test (von Frey test)

Mechanical sensitivity of the right hind paw was assessed using a calibrated series of von Frey hairs (0.02–6□g, Stoelting, Wood Dale, IL, USA) following the up-down method^46,47^. Tests were performed after at least three habituations, each at 24-hour intervals. Assessments were made 1□day before surgery for baseline, and 1, 3, and 7□days after SNT. Rapid paw withdrawal, licking, and flinching were interpreted as signs of pain. All behavioral tests were performed in a blinded manner to the conditions. Before the von Frey test, we confirmed that none of the mice exhibited alterations in other behaviors, including locomotor activity.

### Immunohistochemistry (IHC)

Mice were transcardially perfused with ice-cold 0.1□M phosphate-buffered saline (PBS; pH 7.4) until all blood was removed, followed by perfusion with ice-cold 4% paraformaldehyde in 0.1□M PBS. Whole spinal cord tissue were post-fixed in 4% paraformaldehyde in 0.1□M PBS overnight at 4°C and cryoprotected with 30% sucrose for 3 days. Coronal 60-μm-thick sections were incubated in cryoprotectant at −20°C until immunohistochemical staining was performed. The sections were incubated for 1□h at room temperature in a blocking solution containing 5% normal goat serum (Jackson ImmunoResearch, Bar Harbor, ME, USA), 2% BSA (Sigma-Aldrich), and 0.1% Triton X-100 (Sigma-Aldrich).

Subsequently, the sections were incubated in the blocking solution with rabbit anti-Iba-1 (019-19741, 1:1,000; Wako, Osaka, Japan), mouse anti-glial fibrillary acidic protein (GFAP) (MAB360, 1:1000; Abcam, Cambridge, UK), rabbit anti-GFAP (Z0334, 1:1000; Dako, Santa Clara, CA, USA) antibodies overnight at 4°C. After being washed with 0.1□M PBS containing 0.1% Triton X-100, the sections were incubated in blocking solution for 1□h with FITC-, Cy3- or Cy5-conjugated secondary antibodies (1:200, Jackson ImmunoResearch) at room temperature, washed three times, and then mounted on gelatin-coated glass slides using Vectashield (Vector Laboratories, Inc., Burlingame, CA, USA). Fluorescent images of the mounted sections were obtained using a confocal microscope (LSM800; Carl Zeiss, Jena, Germany).

### qPCR

The quantitative PCR (qPCR) experiments were performed using a StepOnePlus real-time PCR system (Applied Biosystems, Foster City, CA, USA) following the 2−ΔΔCt method. Total RNA from contralateral Spinal Cord tissue was extracted using TRIzol (Invitrogen) and reverse transcribed using TOPscript RT DryMIX (Enzynomics, Cat # RT200, Daejeon, Korea). All the ΔCt values were normalized to the corresponding GAPDH values, and represent fold change induction.

The following qPCR primers were used:

*Gfap* Fw: CAC CTA CAG GAA ATT GCT GGA GG;

*Gfap* Rv: CCA CGA TGT TCC TCT TGA GGT G;

*Iba1* Fw: TCT GCC GTC CAA ACT TGA AGC C;

*Iba1* Rv: CTC TTC AGC TCT AGG TGG GTC T;

*Fasn* Fw: CAC AGT GCT CAA AGG ACA TGC C;

*Fasn* Rv: CAC CAG GTG TAG TGC CTT CCT C;

*Elovl5* Fw: TCG ATG CGT CAC TCG TAC CTA TT;

*Elovl5* Rv: ATT TTG GTC CCA GCC ATA CAA T;

*Elovl6* Fw: CGG CAT CTG ATG AAC AAG CGA G;

*Elovl6* Rv: GTA CAG CAT GTA AGC ACC AGT TC;

*Ldha* Fw: AAA GAG GAC TAA GGG GTG GC;

*Ldha* Rv: CTG CAG GAA ACA ACC ACT CC;

*Hk2* Fw: CCC TGT GAA GAT GTT GCC CAC T;

*Hk2* Rv: CCT TCG CTT GCC ATT ACG CAC G;

*Acod1* Fw: GGT ATC ATT CGG AGG AGC AAG AG;

*Acod1* Rv: ACA GTG CTG GAG GTG TTG GAA C;

### GC-MS

Tissue samples for the GC-MS analysis were prepared from the spinal cord (SC). We collected ipsilateral lateral L3–L5 spinal cord tissue at day 7 post-surgery and performed GC-MS. Samples were homogenized using a tissue homogenizer, followed by heat-based enzyme inactivation at 90 °C for 10 min. The homogenates were then centrifuged at 12,000 rpm at 4 °C for 10 min, and 400 µL of tissue supernatant was collected from each SC specimen for GC-MS analysis.

Derivatization was performed using trimethylsilylation, and 1 µL of each derivatized sample was injected into a Thermo Scientific ISQ LT GC-MS system (Thermo Scientific, Waltham, MA, USA). Chromatographic separation and mass detection were performed under standard operating conditions^48^.

Raw data were processed using Thermo Xcalibur Quan Browser. Metabolites were identified by matching each chromatographic peak to reference spectra, and peak areas were integrated against the baseline to obtain relative signal intensities. All metabolite abundances were then normalized to the internal standard peaks.

### Metabolomics analysis

Absolute and relative metabolite concentrations obtained by GC-MS, including fold-change values normalized to the sham controls, were subjected to a comprehensive metabolomics analysis. For each metabolite, both the absolute abundance and the fold change relative to the sham and SNT groups were calculated, and statistical significance was assessed.

PCA and PLS-DA were performed in MetaboAnalyst using the normalized metabolite abundance matrix, which was generated by sum normalization followed by log_2_transformation, rather than using fold-change values. To identify enriched metabolic pathways within each experimental group, we performed both an over-representation analysis and a quantitative enrichment analysis. All pathway enrichment analyses were conducted using MetaboAnalyst v6.0. (http://www.metaboanalyst.ca)^21,22^.

### RNA-seq data analysis

Raw RNA-seq count matrices were processed in Python for parsing and quality checks, and differential expression between the experimental and control groups was evaluated using DESeq2 with the sham group as the reference. Library-size normalization was performed using DESeq2 size-factor estimation, and gene-wise differential expression statistics (log_2_fold change and associated p-values) were obtained by fitting a negative binomial generalized linear model. For enrichment analyses, genes were pre-ranked by the signed Wald statistic (log_2_FC divided by its standard error). Hallmark pathway enrichment was assessed using the Broad Institute GSEA software (pre-ranked mode) with MSigDB Hallmark gene sets. KEGG pathway enrichment was evaluated using the WEB-based GEne SeT AnaLysis Toolkit (WebGestalt; https://www.webgestalt.org/) using the ranked gene list^49^. Normalized enrichment scores and false discovery rates were computed for each analysis, and enrichment outputs were exported and visualized for figure preparation (GraphPad Prism). Enrichment results were generated for each dataset independently and then compared across datasets to identify pathways showing consistent directional changes^50,51^.

### Quantification and statistical analysis

The data were analyzed in GraphPad software. Student’s *t-*test was used for comparisons between two groups. All data are expressed as the mean ± standard error of the mean (SEM), and statistical significance was defined as a p-value < 0.05.

## Notes

### Competing Interest Statement

The authors have declared no competing interest.

### Summary of Updates

Toning down broad claims about chronic pain, generalizability, lipid metabolism, and biomarker relevance. Revising the title to emphasize reduced free fatty-acid pools. Adding new validation data: Fig. 1C-E, Fig. S1, and Fig. 5. Clarifying the RNA-seq reanalysis and expanding the Methods section. Correcting figure legends, terminology, behavioral time points, references, and PCA/PLS-DA descriptions.

## REFERENCES

1. Colloca L et al (2017) Neuropathic pain. Nat Rev Dis Primers 3:17002.

2. Carlton SM, D. J, Tan HY, Nesic O, Hargett GL, Bopp AC, Yamani A, Lin Q, Willis WD, Hulsebosch CE (2009) Peripheral and central sensitization in remote spinal cord regions contribute to central neuropathic pain after spinal cord injury. Pain 147:265–276.

3. Donnelly CR, Andriessen AS, Chen G, Wang K, Jiang C, Maixner W, Ji RR (2020) Central nervous system targets: glial cell mechanisms in chronic pain. Neurotherapeutics 17:846–860.

4. Mika J, Zychowska M, Popiolek-Barczyk K, Rojewska E, Przewlocka B (2013) Importance of glial activation in neuropathic pain. Eur J Pharmacol 716:106–119.

5. Chen G, Zhang Y, Qadri YJ, Serhan CN, Ji RR (2018) Microglia in pain: detrimental and protective roles. Neuron 100:1292–1314.

6. Ji RR, Donnelly CR, Nedergaard M (2019) Astrocytes in chronic pain and itch. Nat Rev Neurosci 20:667–685.

7. Chen YL, Feng XL, Cheung CW, Liu JA (2022) Mode of action of astrocytes in pain: from the spinal cord to the brain. Prog Neurobiol 102365.

8. Xu Q et al (2021) Astrocytes contribute to pain gating in the spinal cord. Sci Adv 7:eabi6287.

9. Marty-Lombardi S, Lu S, Ambroziak W, Schrenk-Siemens K, Wang J, DePaoli-Roach AA, Hagenston AM, Wende H, Tappe-Theodor A, Simonetti M et al (2024) Neuron-astrocyte metabolic coupling facilitates spinal plasticity and maintenance of inflammatory pain. Nat Metab 6:494–513.

10. Díaz-García CM (2024) Glycogen from spinal astrocytes dials up the pain. Nat Metab 6:384–386.

11. Chen Y, Liao Y, Zhu H et al (2025) Sox9 regulation of hexokinase 1 controls neuroinflammatory astrocyte subtypes in a rat model of neuropathic pain. Nat Commun 16:10249.

12. Park JS, Kim KH, Jun HW, Choi SY, Park SB, Lee SJ (2026) Astrocytic glycogenolysis gates Warburg-like metabolic reprogramming that promotes neuropathic pain chronification. Prog Neurobiol 102924.

13. Fotio Y et al (2021) NAAA-regulated lipid signaling governs the transition from acute to chronic pain. Sci Adv 7:eabi8834.

14. Zhang H, Fan J, Kong D, Sun Y, Zhang Q, Xiang R, Lu S, Yang W, Feng L, Zhang H (2025) Immunometabolism: crosstalk with tumor metabolism and implications for cancer immunotherapy. Mol Cancer 24:249.

15. Lonkar N, Latz E, McManus RM (2025) Neuroinflammation and immunometabolism in neurodegenerative diseases. Curr Opin Neurol 38:163–171.

16. Xu S, Wang Y (2024) Transient receptor potential channels: multiple modulators of peripheral neuropathic pain in several rodent models. Neurochem Res 49:872–886.

17. Kim D, You B, Jo EK, Han SK, Simon MI, Lee SJ (2010) NADPH oxidase 2-derived reactive oxygen species in spinal cord microglia contribute to peripheral nerve injury-induced neuropathic pain. Proc Natl Acad Sci U S A 107:14851–14856.

18. Lee J, Noh K, Lee S et al (2025) Ganglioside GT1b prevents selective spinal synapse removal following peripheral nerve injury. EMBO Rep 26:2994–3023.

19. Lee J, Hwang H, Lee SJ (2021) Distinct roles of GT1b and CSF-1 in microglia activation in nerve injury-induced neuropathic pain. Mol Pain 17.

20. Lim H, Lee J, You B et al (2020) GT1b functions as a novel endogenous agonist of toll-like receptor 2 inducing neuropathic pain. EMBO J 39:EMBJ2019102214.

21. Pang Z, Lu Y, Zhou G, Hui F, Xu L, Viau C, Spigelman AF, MacDonald PE, Wishart DS, Li S, Xia J (2024) MetaboAnalyst 6.0: towards a unified platform for metabolomics data processing, analysis and interpretation. Nucleic Acids Res 52:W398–W406.

22. Ewald JD, Zhou G, Lu Y, Kolic J, Ellis C, Johnson JD, MacDonald PE, Xia J (2024) Web-based multiomics integration using the Analyst software suite. Nat Protoc 19:1467–1497.

23. Wang C, Zhang L, Li P (2024) RNA expression profiling of spinal cord tissues in a rat model of neuropathic pain. Gene Expression Omnibus GSE256472.

24. Cao S, Yuan J, Zhang D, Wen S, Wang J, Li Y, Deng W (2019) Transcriptome changes in dorsal spinal cord of rats with neuropathic pain. J Pain Res 12:3013–3023.

25. Li R, Zhang W, Bai X, Wang F, Yao M, Zhao C, Wang J (2025) Duhuo Jisheng Mixture attenuates neuropathic pain by inhibiting S1PR1/P2Y1R pathway after chronic constriction injury in mice. Phytomedicine 138:156413.

26. Li Y, Yin C, Liu B et al (2021) Transcriptome profiling of long noncoding RNAs and mRNAs in spinal cord of a rat model of paclitaxel-induced peripheral neuropathy identifies potential mechanisms mediating neuroinflammation and pain. J Neuroinflammation 18:48.

27. Currie E, Schulze A, Zechner R, Walther TC, Farese RV Jr (2013) Cellular fatty acid metabolism and cancer. Cell Metab 18:153–161.

28. Wang X, Yu H, Gao R, Liu M, Xie W (2023) A comprehensive review of the family of very-long-chain fatty acid elongases: structure, function, and implications in physiology and pathology. Eur J Med Res 28:532.

29. Tannahill G, Curtis A, Adamik J et al (2013) Succinate is an inflammatory signal that induces IL-1β through HIF-1α. Nature 496:238–242.

30. Mills E, Ryan D, Prag H et al (2018) Itaconate is an anti-inflammatory metabolite that activates Nrf2 via alkylation of KEAP1. Nature 556:113–117.

31. Llibre A, Kucuk S, Gope A, Certo M, Mauro C (2025) Lactate: a key regulator of the immune response. Immunity 58:535–554.

32. Lampropoulou V, Sergushichev A, Bambouskova M et al (2016) Itaconate links inhibition of succinate dehydrogenase with macrophage metabolic remodeling and regulation of inflammation. Cell Metab 24:158–166.

33. Lang R, Siddique MNAA (2024) Control of immune cell signaling by the immuno-metabolite itaconate. Front Immunol 15:1352165.

34. Liu Y, Jiang X, Zhuang S, Zhu L, Zhu B, Rui K, Tian J (2025) Itaconate: a key regulator of immune responses and potential therapeutic target for autoimmune and inflammatory diseases. Autoimmun Rev 24:103885.

35. Aguayo-Cerón KA, Sánchez-Muñoz F, Gutierrez-Rojas RA, Acevedo-Villavicencio LN, Flores-Zarate AV, Huang F, Giacoman-Martinez A, Villafaña S, Romero-Nava R (2023) Glycine: the smallest anti-inflammatory micronutrient. Int J Mol Sci 24:11236.

36. Georgieva M, Wei Y, Dumitrascuta M, Pertwee R, Finnerup NB, Huang W (2019) Fatty acid suppression of glial activation prevents central neuropathic pain after spinal cord injury. Pain 160:2724–2742.

37. Yan J, Horng T (2020) Lipid metabolism in regulation of macrophage functions. Trends Cell Biol 30:979–989.

38. Buck MD, Sowell RT, Kaech SM, Pearce EL (2017) Metabolic instruction of immunity. Cell 169:570–586.

39. Chen YY, Feng LM, Xu DQ, Yue SJ, Fu RJ, Zhang MM, Tang YP (2022) Combination of paeoniflorin and liquiritin alleviates neuropathic pain by lipid metabolism and calcium signaling coordination. Front Pharmacol 13:944386.

40. Banno T, Omura T, Masaki N, Arima H, Xu D et al (2017) Arachidonic acid-containing phosphatidylcholine increases due to microglial activation in ipsilateral spinal dorsal horn following spared sciatic nerve injury. PLoS One 12:e0177595.

41. Yekkirala AS, Roberson DP, Bean BP, Woolf CJ (2017) Breaking barriers to novel analgesic drug development. Nat Rev Drug Discov 16:545–564.

42. Ji RR, Xu ZZ, Gao YJ (2014) Emerging targets in neuroinflammation-driven chronic pain. Nat Rev Drug Discov 13:533–548.

43. Mabou Tagne A, Fotio Y, Lee HL, Jung KM, Katz J, Ahmed F, Le J, Bazinet R, Jang C, Piomelli D (2025) Metabolic reprogramming in the spinal cord drives the transition to pain chronicity. Cell Rep 44:116261.

44. Park JS, Lee DG, Min DH, Lee SJ (2025) Glioblastoma immunotherapy adjuvants for glial cell polarization regulation. Exp Neurobiol 34:235–247.

45. Pålsson-McDermott EM, O’Neill LAJ (2020) Targeting immunometabolism as an anti-inflammatory strategy. Cell Res 30:300–314.

46. Tanga FY, Nutile-McMenemy N, DeLeo JA (2005) The CNS role of toll-like receptor 4 in innate neuroimmunity and painful neuropathy. Proc Natl Acad Sci U S A 102:5856–5861.

47. Chaplan SR, Bach FW, Pogrel JW, Chung JM, Yaksh TL (1994) Quantitative assessment of tactile allodynia in the rat paw. J Neurosci Methods 53:55–63.

48. Chan E, Pasikanti K, Nicholson J (2011) Global urinary metabolic profiling procedures using gas chromatography–mass spectrometry. Nat Protoc 6:1483–1499.

49. Wang J, Vasaikar S, Shi Z, Greer M, Zhang B (2017) WebGestalt 2017: a more comprehensive, powerful, flexible and interactive gene set enrichment analysis toolkit. Nucleic Acids Res 45:W130–W137.

50. Liao Y, Wang J, Jaehnig EJ, Shi Z, Zhang B (2019) WebGestalt 2019: gene set analysis toolkit with revamped UIs and APIs. Nucleic Acids Res 47:W199–W205.

51. Elizarraras JM, Liao Y, Shi Z, Zhu Q, Pico AR, Zhang B (2024) WebGestalt 2024: faster gene set analysis and new support for metabolomics and multi-omics. Nucleic Acids Res 52:W415–W421.

