## Supplementary Information for "Peripheral nerve injury induces an immunometabolic signature involving reduced free fatty-acid pools"

| **Contents** | **Page** |
|---|---|
| Supplementary Figures | 2 |

### Supplementary Figures

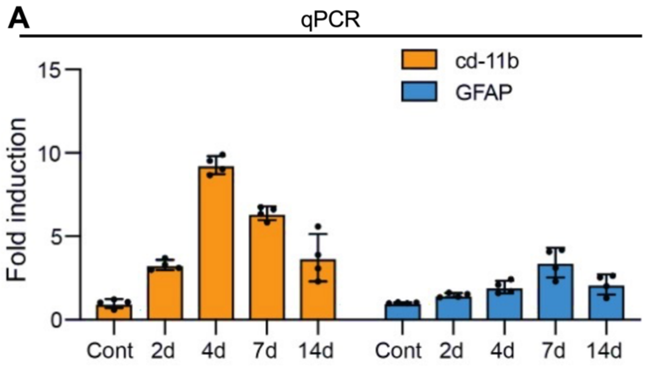


**Figure S1. Time-dependent induction of microglial and astrocytic markers after injury**

(A) qPCR analysis showing temporal changes in cd-11b and GFAP expression following injury. cd-11b expression increased prominently at 4 d and 7 d, whereas GFAP showed a later and moderate increase, peaking around 7 d. Data are presented as fold induction relative to control.

Bars indicate mean ± SEM.

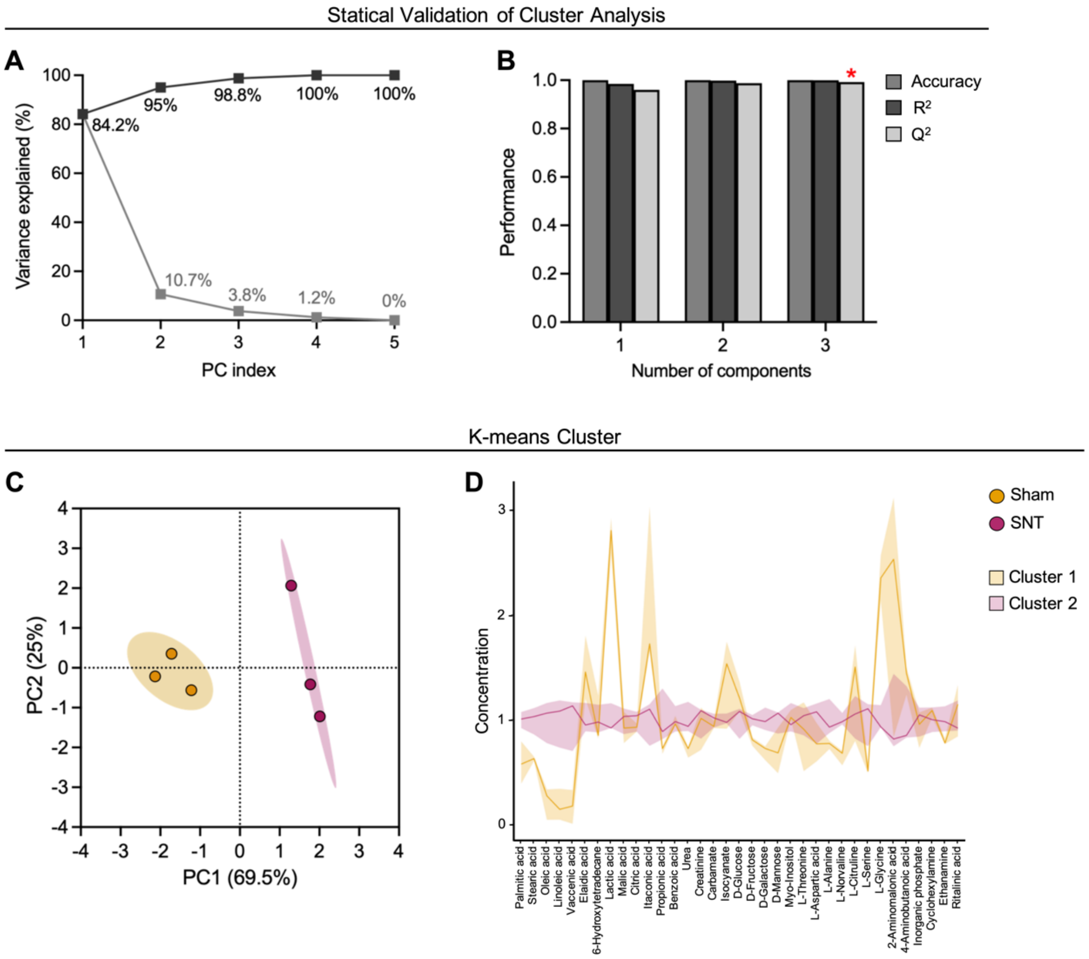


**Figure S2. Statistical validation of cluster separation in GC–MS metabolomics. (Related in Fig 2.)**

(A) Variance explained by principal components (PCs) from PCA of GC–MS metabolite profiles. The percentage of variance explained by each PC (gray) and cumulative variance explained (black) are shown.

(B) Model performance metrics across the number of components, including accuracy, $R^2$, and $Q^2$ (as indicated in the legend).

(C) PCA score plot showing separation between sham and SNT groups with confidence ellipses; each point represents an individual sample.

(D) K-means clustering results visualized as coefficient profiles across metabolites, with group-level trends shown for sham vs SNT and Cluster 1 vs Cluster 2 (as indicated in the legend).

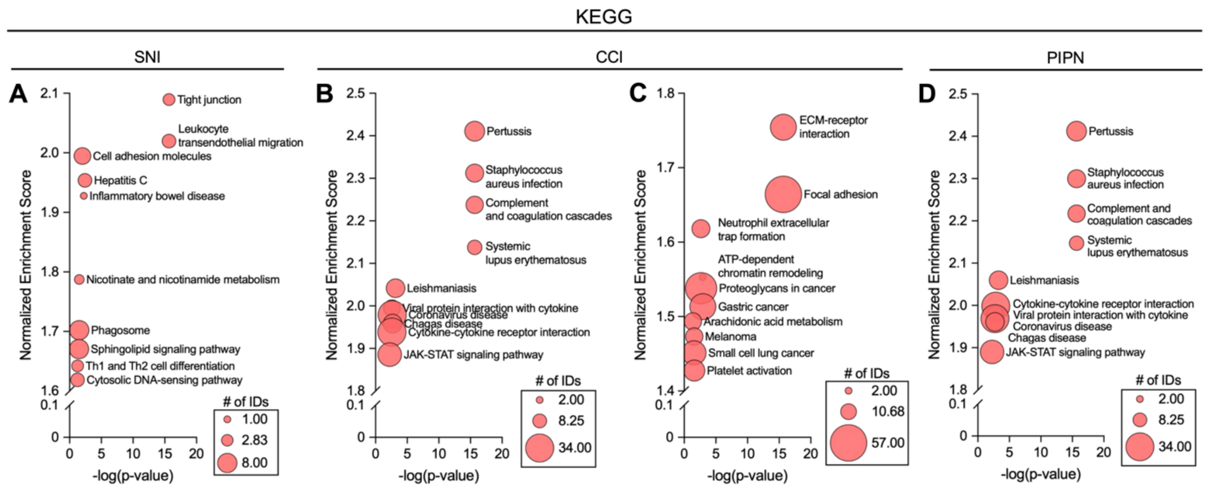


**Figure S3. KEGG pathway enrichment of upregulated programs across independent spinal cord RNA-seq datasets**

KEGG pathway enrichment results (WebGestalt) for genes showing positive enrichment in each neuropathic pain dataset. Bubble plots display normalized enrichment score (NES) on the y-axis and −log(p-value) on the x-axis; bubble size reflects the number of mapped gene IDs (# of IDs).

(A–D) Enriched KEGG pathways in SNI (A), CCI (B), CCI2 (C), and PIPN (D) datasets.

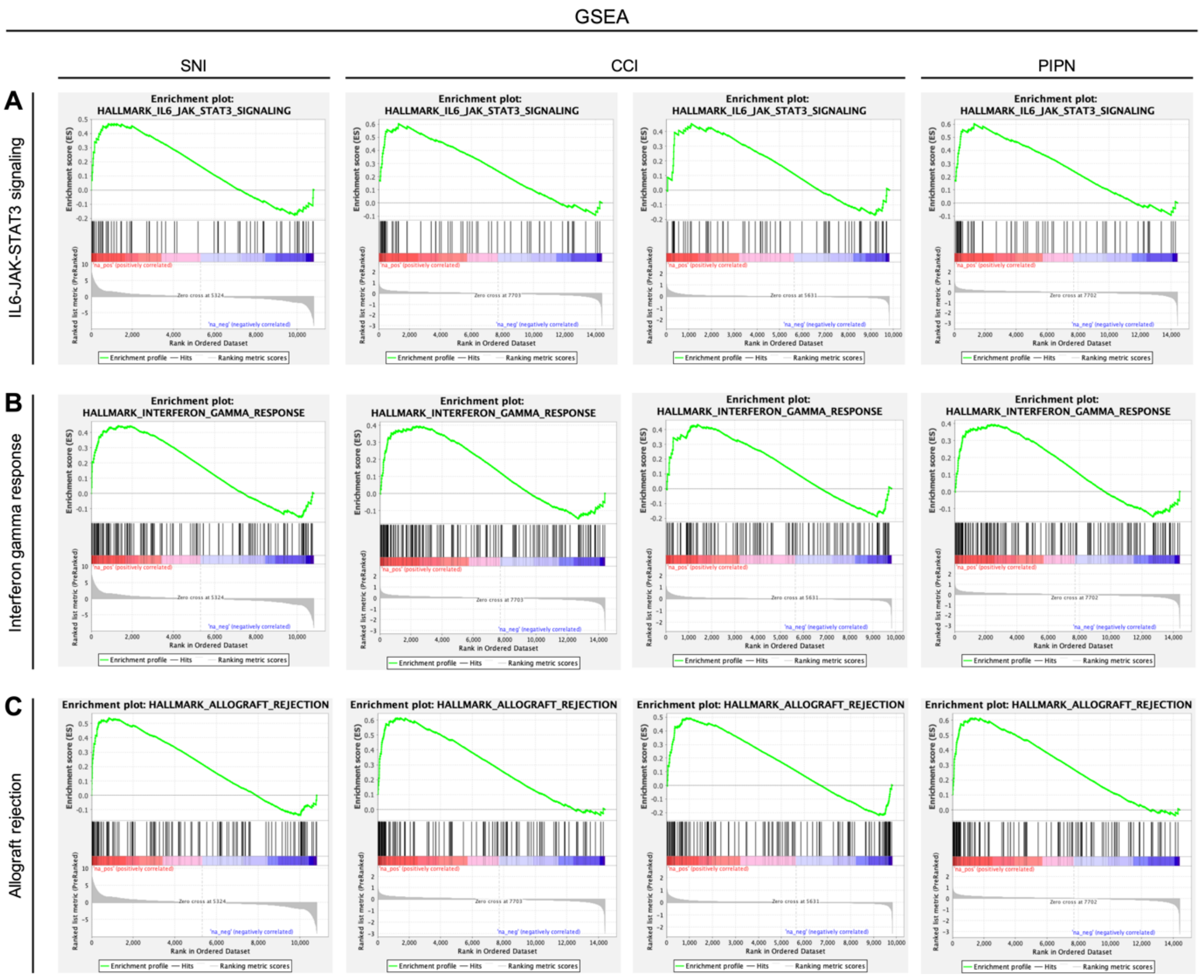


**Figure S4. Hallmark GSEA enrichment plots for inflammatory/immune-related gene sets across datasets**

Representative Hallmark gene-set enrichment plots (GSEA, pre-ranked mode) across four independent spinal cord RNA-seq datasets. Columns correspond to SNI, CCI, CCI2, and PIPN.

(A) HALLMARK_IL6_JAK_STAT3_SIGNALING.

(B) HALLMARK_INTERFERON_GAMMA_RESPONSE.

(C) HALLMARK_ALLOGRAFT_REJECTION.

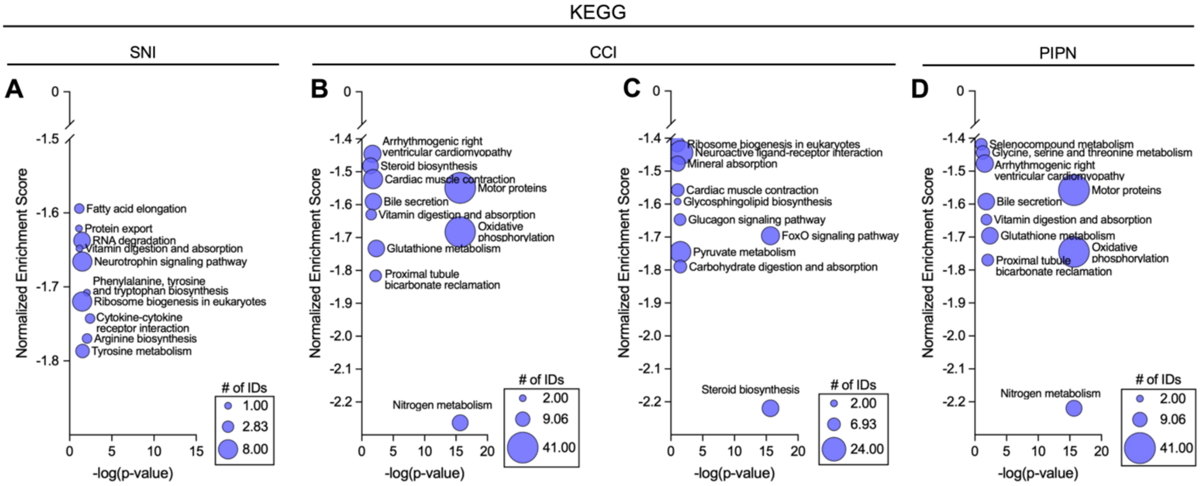


**Figure S5. KEGG pathway enrichment of downregulated metabolic programs across independent RNA-seq datasets**

KEGG pathway enrichment results (WebGestalt) for genes showing negative enrichment in each dataset. Bubble plots display NES (negative values) on the y-axis and −log(p-value) on the x-axis; bubble size reflects the number of mapped gene IDs (# of IDs).

(A–D) Enriched KEGG pathways in SNI (A), CCI (B), CCI2 (C), and PIPN (D) datasets.

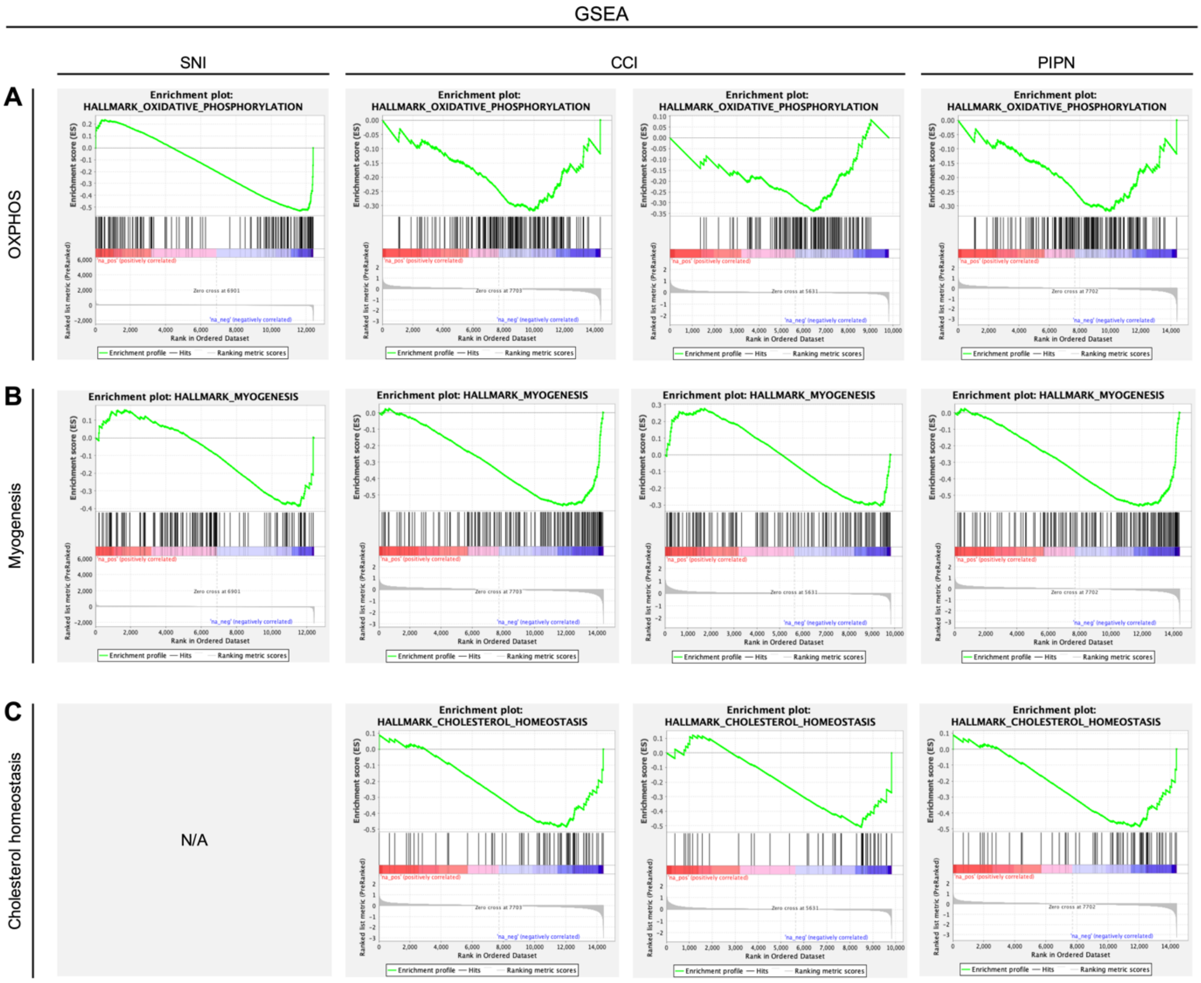


**Figure S6. Hallmark GSEA enrichment plots for metabolic programs across datasets**

Representative Hallmark gene-set enrichment plots (GSEA, pre-ranked mode) across four independent spinal cord RNA-seq datasets. Columns correspond to SNI, CCI, CCI2, and PIPN.

(A) HALLMARK_OXIDATIVE_PHOSPHORYLATION.

(B) HALLMARK_MYOGENESIS.

(C) HALLMARK_CHOLESTEROL_HOMEOSTASIS (N/A indicates that the plot was not available for the corresponding dataset, as shown).
